# Urbanization is associated with reduced genetic diversity in marine fish populations

**DOI:** 10.1101/2024.02.20.581210

**Authors:** Eleana Karachaliou, Chloé Schmidt, Evelien de Greef, Margaret F Docker, Colin J Garroway

## Abstract

The economic and ecological benefits of living by the ocean have led many coastal settlements to grow into large densely populated cities. Large coastal cities have had considerable environmental effects on marine ecosystems through resource extraction, waste disposal, and use for transportation. Thus, it is important to understand the consequences of urbanization and human activities on evolutionary processes and biodiversity in marine fishes. Using published population genetic datasets for marine fishes amounting to 75,496 individuals sampled from 73 species at 1143 sample sites throughout the world’s oceans, we evaluated how human population density and a composite measure of cumulative human impacts affected genetic diversity and differentiation. We found that genetic diversity was significantly lower in marine fish populations associated with denser human populations regardless of species and locality. The effects of cumulative human impacts on genetic diversity were less prominent, perhaps due to this measure capturing more spatially varying processes. Urbanization in coastal regions has degraded marine biodiversity in a way that erodes adaptive potential for marine fish populations. This highlights the need to mitigate threats from human activities and focus efforts on sustainable urban planning and resource use to conserve marine biodiversity sustaining coastal fisheries and ecosystems.

## Introduction

Throughout history, oceans have provided humans with substantial food resources and a means for waste disposal^1–3^ and transporting goods and people^4^. This has transformed many coastal cities into regional and global centers of wealth and population growth^5^. Today, more than 40% of the human population lives in densely populated cities within 100 km of the ocean^6^. The environmental footprint of coastal cities has grown with their economies^5^, increasing oceanic habitat conversion, pollution, and resource exploitation^7–9^. Intact coastal regions are now globally rare—approximately 15% of coasts are considered minimally pressured by human settlement and use^10^. Despite the long history of coastal settlements and the extended reach of urbanization into marine environments, we know little about how urbanization has affected evolutionary processes and biodiversity in the oceans^11,12^. We hypothesized that human activities in coastal cities have reduced the capacity for marine habitats near cities to support large, intact fish populations, resulting in population decline and loss of genetic diversity.

Declines and collapses of exploited marine fish populations due to overfishing have had well-documented and significant negative ecological and socioeconomic consequences^13–15^. However, global threats to marine fish biodiversity at the genetic level are underexplored (but see ^16^ for the genetic consequences of fishing), particularly for species that are not commercially harvested. Genetic diversity underlies all facets of biodiversity, including population persistence, the capacity to adapt to environmental change^17^ and, ultimately, ecosystem stability and resilience^18^. The erosion of genetic diversity is thus a significant ecological concern for maintaining functioning marine ecosystems and sustainable fisheries. However, assessments of genetic diversity are not routinely conducted as part of population monitoring and commercial harvest of marine fishes. While our understanding of the natural drivers of genetic variation is beginning to be explored^19,20^, we lack basic information about the threats posed to marine fish genetic diversity by human settlements.

We took advantage of the accumulation of raw population genetic datasets archived with individual research papers to test whether fish populations near big cities tended to have reduced genetic diversity for marine fish species at a global scale. We focused on two measures of human disturbance: human population density^21^ and a composite measure of cumulative human impacts^22^ which includes effects of settlements together with estimates of the effects of climate change, fishing pressure, and numerous other anthropogenic threats to the marine environment. Human population density should capture the degree of urbanization and the associated long-term, spatially consistent effects of humans on natural environments. While the nature of the effects of humans on environments will vary locally, we hypothesized that they will generally be negative. We predicted that fish population size, and thus genetic diversity, would decrease, and populations would become increasingly genetically differentiated with increasing exposure to urbanization and cumulative environmental effects of humans.

## Results and Discussion

Our full data set comprised 75,496 individuals sampled from 73 marine fish species, including diadromous and brackish species, at 1143 sample sites throughout the world’s oceans (Fig 1). Allowing the relationship between the genetic composition of populations and urbanization to vary with species, we found a globally coherent signature of the erosion of genetic diversity in marine fish populations associated with denser human populations.

**Figure 1.**
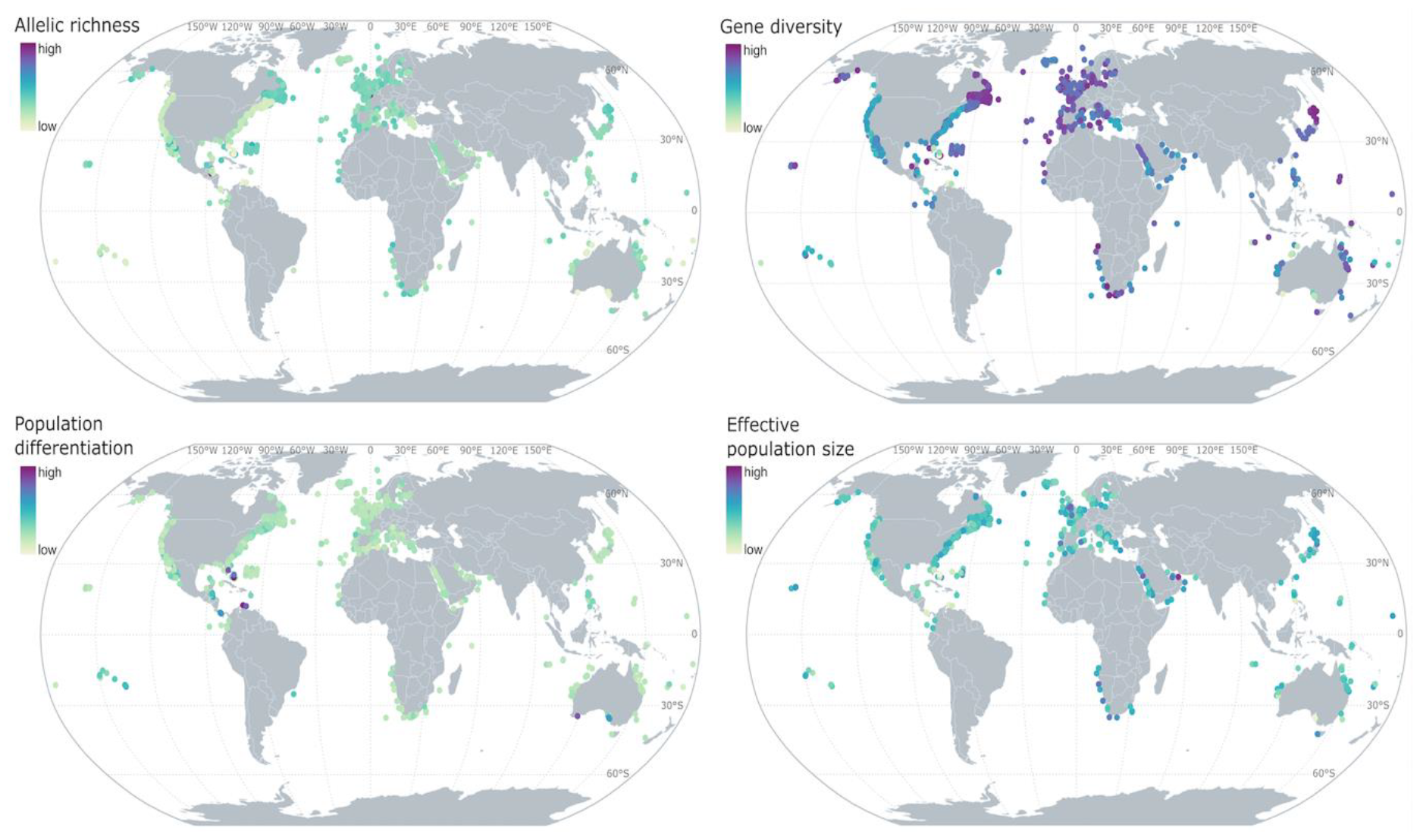
Global maps of per population raw gene diversity, allelic richness, population differentiation (FST), and effective population size. Each point on the map is a sample site where a single species was sampled.

Populations associated with dense human settlements had reduced allelic richness, gene diversity, and effective population size (Table 1; Fig 2). The consistent direction of effects suggests these relationships hold generally across taxa and sample locality. There was no detectable relationship between human population density and genetic differentiation.

**Table 1.** Model summaries for the effects of human population density and cumulative human impacts on genetic diversity and differentiation. Mean overall effect sizes for each predictor are given with 95% credible intervals. The widely applicable information criterion (WAIC) is an indicator of relative model fit. Probability of direction (pd) is a measure of the probability an effect is positive or negative. R^2^ values represent coefficient of variation and indicate model fit. Bayesian R^2^ is the variance of the predicted values divided by the variance of predicted values plus the expected variance of the errors.

| | predictor | mean (95% CI) | pd | Bayesian $R^2$ | WAIC |
| --- | --- | --- | --- | --- | --- |
| gene diversity | human population density | -0.10 (-0.17 -0.01) | 99% | 0.83 | 1173.8 |
|  | cumulative human impacts | -0.04 (-0.11 0.03) | 88% | 0.81 | 1374.1 |
| allelic richness | human population density | -0.06 (-0.15 0.02) | 95% | 0.75 | 1587.1 |
|  | cumulative human impacts | -0.06 (-0.13 0.00) | 97% | 0.75 | 1661.1 |
| population-specific $F_{ST}$ | human population density | 0.04 (-0.12 0.21) | 70% | 0.55 | 2198.8 |
|  | cumulative human impacts | 0.02 (-0.08 0.12) | 66% | 0.49 | 2408.7 |
| effective population size | human population density | -0.09 (-0.23 0.03) | 94% | 0.28 | 2412.4 |
|  | cumulative human impacts | 0.05 (-0.05 0.17) | 82% | 0.32 | 1733.7 |

**Figure 2.**
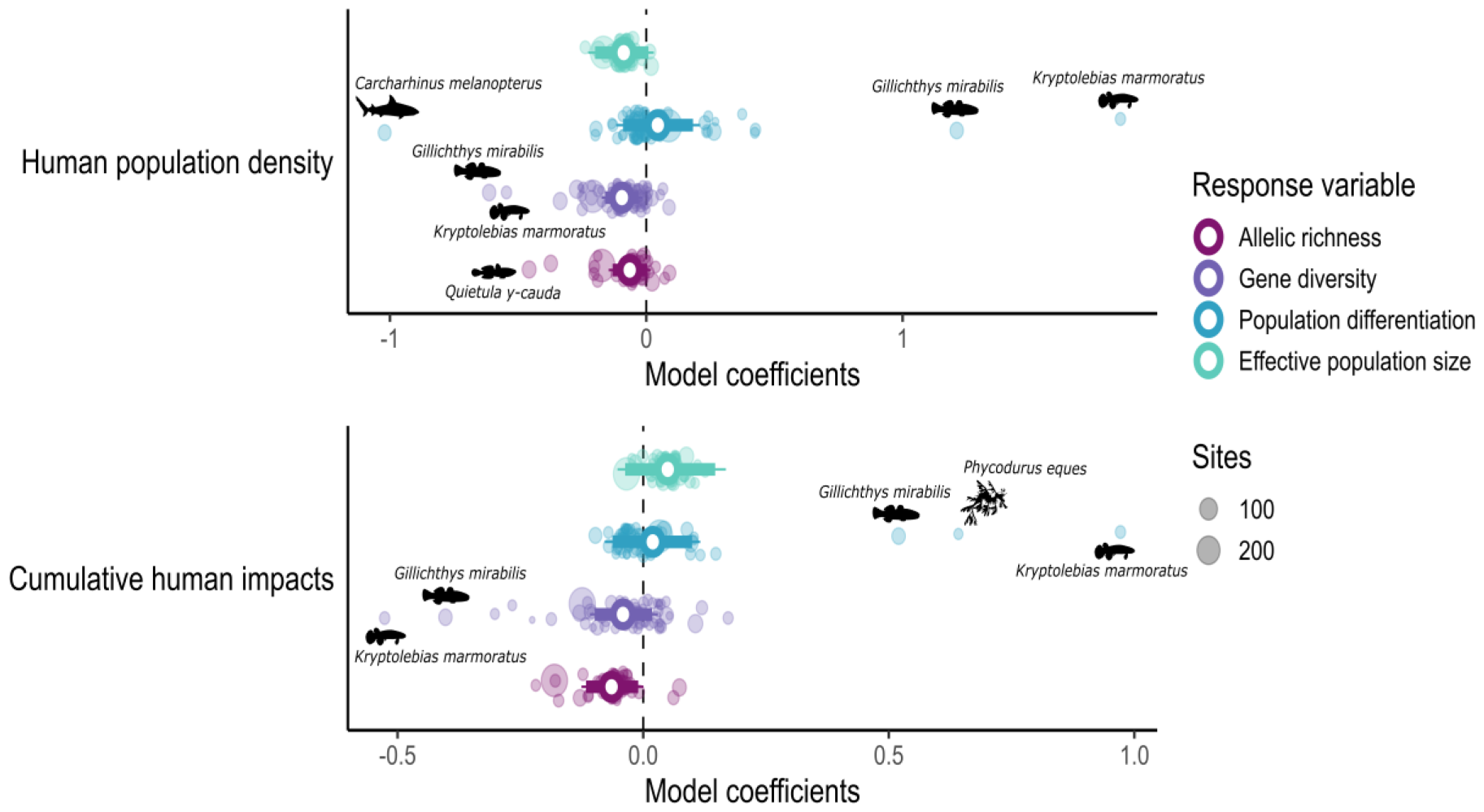
The results of generalized linear mixed effects models for the relationship between human population density and cumulative human impacts and allelic richness, gene diversity, population differentiation, and effective population size. We allowed slopes and intercepts to vary with species. Open circles are overall parameter estimates, bold lines are 90% credible intervals and narrow lines are 95% credible intervals. Faded points are species specific parameter estimates with outliers indicated by species name and silhouette.

Relationships between cumulative human impacts and genetic diversity and differentiation were less clear (Table 1; Fig 2). Allelic richness and gene diversity were both negatively associated with cumulative human impacts, but relationships were weaker than those between human population density and the genetics of sample sites. There was also little evidence for a relationship between cumulative human impacts and population genetic differentiation.

Unlike associations with human population density, we found no detectable relationship between cumulative human impacts and effective population size. These weaker associations between cumulative human impacts and the genetic composition of populations may be due to the inclusion of spatiotemporally varying factors in the composite index. For example, variations in fishing pressure and sea surface temperature anomalies vary across space and time, contrasting with the typically spatially fixed and persistent effects of human population densities in coastal cities, which tend to increase over time. The lack of relationship between urbanization and genetic differentiation suggests that reductions in genetic diversity are due to the reduced capacity to support fish populations rather than habit fragmentation.

Reduced genetic diversity in association with urbanization suggests adaptive capacity is reduced in marine fish populations near cities. Maintaining genome-wide genetic variation prevents inbreeding depression and maintains adaptive potential^23^. Further, the efficiency of natural selection in a population is inversely proportional to its effective population size. As effective population size declines, the random process of genetic drift can come to overpower the more deterministic process of natural selection, with even well-adapted alleles increasingly being lost to chance during Mendelian segregation. Our findings thus suggest that coastal marine fish populations and ecosystems are less resilient to future environmental change and continued exploitation and urban expansion.

The cumulative human impacts index included fishing pressure, a direct contributor to population decline, and so we had predicted it would have a more detectable signature of genetic erosion. Previous meta-analytic work exploring the genetic consequences of overfishing found that fishing reduced genetic diversity^16^. The weak associations we found between the index of cumulative human impacts and the genetic composition of populations do not necessarily contradict this earlier result. Our analyses were concerned with declines of genetic diversity across populations within species, whereas Pinsky and Palumbi^16^ addressed this question across species using sister taxon comparisons between overfished species and species not known to be overfished. Taken together, these two analyses suggest that fished species lose genetic diversity on the whole^16^, with little evidence for population-level variation in the genetic consequences of fishing pressure. If true, genetic rescue, the movement of alleles from one population to another that increases genetic diversity, should not generally be expected to occur and may not be a viable conservation tool in exploited marine fish populations.

We are moving toward an increasingly complete picture of ways that urbanization shapes neutral genetic diversity in wild populations. Human activities are now the principal driver of evolutionary change in wild species^24,25^. When we apply selection pressures on populations directly, such as through pest control^26^, evolutionary change is predictable—we can expect pest populations to evolve adaptively in response to intentionally applied selection pressures^23^. However, where habitat disruption is more haphazard, as it is with urbanization, evolutionary responses can be idiosyncratic. On land, urbanization reduces mammalian genetic diversity and increases genetic isolation^27^. Avian genetic diversity was also consistently affected by urbanization, but effects varied between positive and negative depending on the species^27^. There are relatively few detectable trends between urbanization and genetic diversity in amphibians, suggesting that the effects of urbanization may not be generalizable for this group^28^. Our findings suggest that when considering the conservation and management of species where the effects of urbanization are unknown, we should assume that they are negative for the genetic diversity of marine fishes. The widespread and extensive urbanization of coastal regions poses threats to marine fish populations, biodiversity, and ecosystems.

Human-caused biodiversity declines are concerning because they threaten the functioning ecosystems on which humans and other species depend. More than one billion people rely on the ocean for their primary food sources^29–31^, and approximately 260 million people rely on fisheries for at least part of their livelihood^32^. Eighty-five percent of marine fisheries are coastal^33^, connected to coastal human communities and population centers.

Relative to developed countries, fisheries play an outsized role in the economy and diet quality of developing countries^13^. The discharge of pollutants, destruction of critical habitats, and unsustainable fishing practices all contribute to the degradation of marine ecosystems^7–9^. Mitigating these threats requires sustainable urban planning, improved wastewater treatment systems, and the implementation of responsible fishing practices. By integrating conservation efforts into urban development, we can protect marine biodiversity and ensure the long-term health and resilience of our oceans.

## Acknowledgements

This work was supported by NSERC Discovery Grants awarded to CJG and MFD and a University of Manitoba Research awarded to MFD. We thank C. Müller of the Population Ecology & Evolutionary Genetics Group at the University of Manitoba for his contribution to the project and D.R. Bath, L. Kathan, K. King, C. Kucheravy, L. Newediuk, R. Rivkin, M. Sherritt, J. Suurväli and J. Tepker of the Population Ecology & Evolutionary Genetics Group at the University of Manitoba for comments on the manuscript. We thank all the authors who made their data publicly available.

## Methods

### Data compilation

We built a database of population genetic datasets for marine fishes following previous work^27,28^. Briefly, we programmatically searched online public data repositories using the DataONE package^34^ in R^35^. We searched species names (e.g., “*Petromyzon marinus*”) and keywords “microsatellite” and “microsat*”. We focused our search on marine fishes, including diadromous and brackish species inhabiting transitional zones. Our species list came from the IUCN Red List database. We chose 1990 as a cut-off for earliest year of data inclusion. We used only presumed neutral microsatellite data, and excluded data from hatchery-bred populations, invasive species, hybrids where noted, and experimental data. We used microsatellites because they are the most commonly used and archived neutral marker type^36^ and because, when sampled at the site level, they are well correlated with genome-wide diversity^37^. We used sample site coordinates when provided in publications and georeferenced^38^ sample sites from maps when necessary. After applying our dataset retention criteria (see SI Fig 1), we were left with 108 microsatellite datasets spanning 73 species, 1143 sample sites, and 75,496 individuals.

We imported the genotype datasets into R using the adegenet package (version 2.1.3^39,40^). We then estimated gene diversity, allelic richness, population-specific FST, and effective population size for each sample site. Gene diversity is a measure of the average probability that two alleles sampled from the population at random are different^41^. We estimated gene diversity with the *Hs*() function in adegenet^39,40^. Allelic richness, an estimate of the number of alleles at a sample site corrected for sample size = 10, was calculated using the *allelic*.*richness()* function in hierfstat^42^. Population-specific FST^43^, a measure of the relative degree of divergence from a common ancestral population, was calculated using the *betas*() function in hierfstat. FST could only be calculated for species with at least 2 sample locations (n = 1131 sites). Contemporary effective population size, the estimated rate at which a population loses genetic diversity due to genetic drift, was estimated for the parental generation based on a linkage disequilibrium method using NeEstimator v2 with a minor allele frequency filter of 0.1^44^. Effective population size was log-transformed for use in further analysis. Estimating effective population is difficult when the sampling error overrides the signal of genetic drift. Effective population size was estimable for 682 sites.

Finally, we filtered out sites with fewer than 5 individuals, leaving 1085 sites, 75,361 individuals, and 73 species for subsequent analysis (SI Table 1).

Repurposing raw data originally generated for different purposes adds robustness to our analyses. This is because we could calculate all population genetic parameters, measures of urbanization, and human impacts consistently and comparably. The datasets we repurposed were collected for the purpose of answering different questions, so it is unlikely that study site, study system, or publication bias affects our findings.

### Urbanization and human impacts at sample sites

We used human population density as an indicator of urbanization and correlated negative effects on associated marine ecosystems. We downloaded estimates of human population density^21^ (number of persons per square kilometer) from national censuses and population registers for the year 2010 in a 30 arc-second resolution. To estimate the sum of human impacts on the world’s oceans we used the index of cumulative human impacts for the year 2013^22^, an indicator of the influence of climate change, fishing, ocean and land-based stressors on marine ecosystems. To test for effects across spatial scales we summarized human population density and human impact variables within buffers of 25, 50, 100 and 200 km radii around each sample site. All datasets were re-projected to the Lambert Cylindrical Equal Area coordinate system to ensure their compatibility and comparability.

### Statistical Analysis

We modelled the relationships between genetic measures, urbanization and human impacts in separate models using Bayesian generalized linear mixed effects models fit in brms^45^.

Predictor and response variables were scaled and centred prior to running models so that effect sizes were comparable across models. We treated species as a random effect and accounted for interspecific variation in the means of each genetic metric with random intercepts, and relationships with predictors by allowing slopes to vary with species. We ran hierarchical models for each measure of genetic diversity as explained by each driver. We ran all models with four chains and a minimum of 2000 iterations. In total we ran 32 models: 4 genetic diversity measures x 2 candidate drivers x 4 buffer sizes. Models were run both with and without priors. We used a conservative normal prior of zero with a standard deviation of one for all fixed effect slope parameter estimates presented in the main text. We tested for spatial autocorrelation in our models using the Moran’s I test in the package spdep^46^. No spatial autocorrelation was present in our model residuals. We estimated the probability of direction of effect for each predictor using the function “p_direction” in bayestestR^47^. This probability measures the proportion of values in the posterior distribution that have the same sign as the median value. Additional model diagnostics were estimated in the brms package^45^ by computing Bayesian R-squared values using the function bayes_R2(), widely applicable information criterion (WAIC) using the function waic() and performing posterior predictive checks using the function pp_check. We interpret models with priors at the most significant buffer per model.

## Notes

### Competing Interest Statement

The authors have declared no competing interest.

