## Supplementary Information for "Urbanization is associated with reduced genetic diversity in marine fish populations"

SI Figure 1. We compiled data in three separate phases, searching the North Atlantic on 10 December 2019 (164 search results, of which 36 were usable), the entire Atlantic on 13 January 2021 (178 search results, of which 44 were usable), and performing a global search on 13 January 2021 (186 search results, of which 59 were usable). Data was considered usable when it consisted of presumed neutral microsatellites (e.g. excluding known adaptive markers and SNPs) generated in a way that would not affect genetic diversity (e.g. excluding individuals reared in captivity and simulated data). There were instances where the same dataset appeared in more than one round of search results. There were also instances where a dataset that was for example found as part of our North Atlantic search but corresponded to the Pacific and was initially dismissed, was later included once the study was extended to include global oceans. After accounting for overlap and including datasets that were initially dismissed due to a mismatch of locality, we retained 84 search results for our global study of the classes Actinopterygii, Chondrichthyes and Petromyzontida.

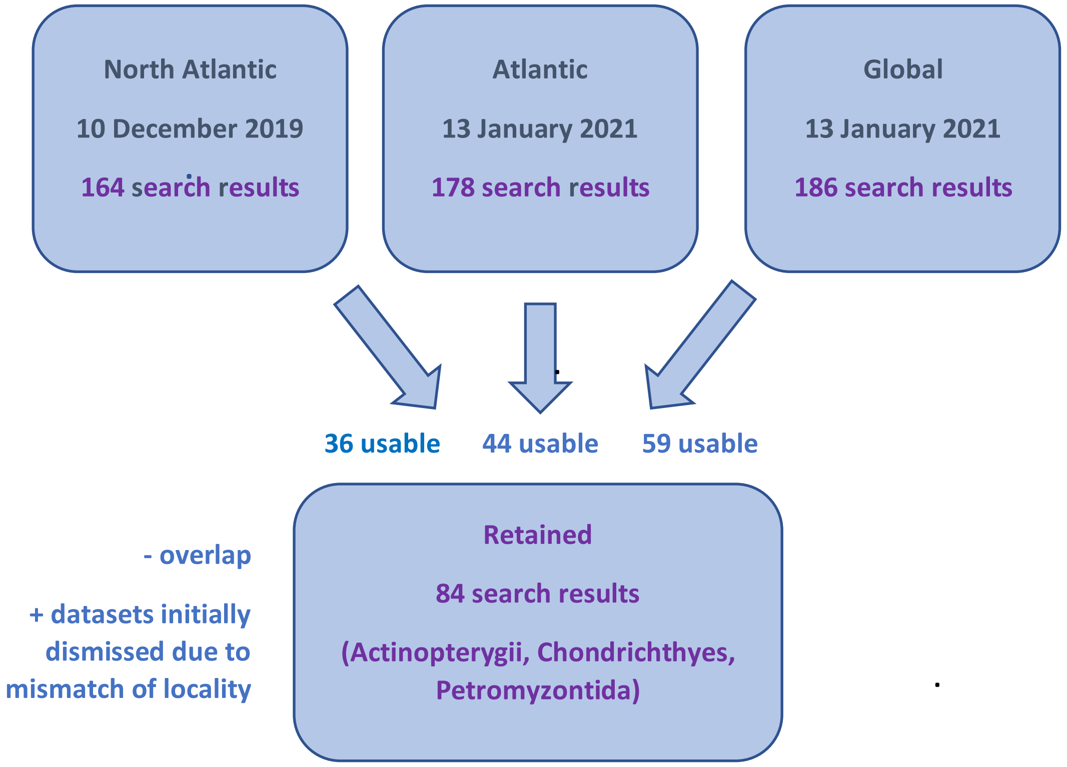


SI Table 1. Summary and reference information for data used in analysis as part of this study. Data was accessed from the Dryad repository. Class: Chondrichthyes (C), Actinopterygii (A) or Petromyzontida (P). Loci: number of microsatellite loci sampled, populations: number of populations, individuals: total number of individuals summed over all populations. Coordinates: method of assigning coordinate locations to each site. C = site coordinates provided in study; AC = coordinates given per sample in study, averaged to obtain site coordinates or neighbouring populations with too few individuals merged so they can be usable, coordinates averaged; GM = site name searched in Google Maps, for anadromous species sampled in freshwater closest marine site to river mouth from Google Maps used; AU

= Author contacted and provided coordinates; M = map provided in study, georeferenced in QGIS; T = Coordinates from northernmost site in transect and per sample distance from northernmost site given in km, coordinates calculated and averaged. Coordinate averages calculated using the geomean() command from the geosphere R package[1]. Further population specific details may apply and are included in the global master population sheet provided in supporting data. Size: Max. length (Lmax: cm) as a measure of body size obtained through the 'rFishbase' package[2]. Exploited: species exploitation status as seen in FishBase (www.fishbase.org). Y= commercially or highly commercially fished species; M = minor commercial fisheries; N = species not commercially fished. Information on species exploitation is for reference only and does not reflect exploitation at the population level. *Accepted species name noted in table, the name used by data authors "*Gobiusculus flavescens*" used in dataset.

| Species | Class | Populations | Coordinates | Individuals | Loci | Search date | Size | Exploited | Reference |
|---|---|---|---|---|---|---|---|---|---|
| *Entosphenus tridentatus* | P | 20 | C, GM | 965 | 9 | 2021 Jan | 76 | N | 3 |
| *Aetobatus narinari* | C | 3 | M | 572 | 5 | 2021 Jan | 230 | M | 4 |
| *Alopias pelagicus* | C | 6 | M | 326 | 7 | 2021 Jan | 428 | Y | 5 |
| *Carcharhinus amblyrhynchos* | C | 9 | C | 112 | 16 | 2021 Jan | 255 | M | 6 |
| *Carcharhinus isodon* | C | 6 | M | 369 | 16 | 2021 Jan | 200 | M | 7 |
| *Carcharhinus limbatus* | C | 10 | M | 812 | 22 | 2021 Jan | 286 | Y | 8,9 |
| *Carcharhinus melanopterus* | C | 31 | M | 1388 | 31 | 2021 Jan | 200 | Y | 10,11 |
| *Carcharhinus sorrah* | C | 6 | M | 700 | 23 | 2021 Jan | 160 | M | 8,9 |
| *Carcharodon carcharias* | C | 2 | M | 166 | 14 | 2019 Dec | 640 | M | 12 |
| *Centroscymnus coelolepis* | C | 8 | C | 503 | 11 | 2019 Dec | 121 | M | 13 |
| *Galeocerdo cuvier* | C | 10 | M | 380 | 10 | 2019 Dec | 750 | Y | 14 |
| *Prionace glauca* | C | 7 | C | 226 | 9 | 2019 Dec | 400 | M | 15 |
| *Rhincodon typus* | C | 7 | M | 406 | 14 | 2021 Jan | 1700 | Y | 16 |
| *Rhizoprionodon acutus* | C | 2 | M | 294 | 8 | 2021 Jan | 175 | Y | 9 |
| *Scyliorhinus canicula* | C | 10 | AC | 834 | 12 | 2019 Dec | 100 | M | 17 |
| *Sphyrna lewini* | C | 5 | M, GM | 342 | 27 | 2021 Jan | 430 | Y | 9,18 |
| *Alosa aestivalis* | A | 28 | GM | 1549 | 15 | 2019 Dec | 40 | Y | 19 |

| | | | | | | | | | |
|---|---|---|---|---|---|---|---|---|---|
| *Alosa pseudoharengus* | A | 56 | GM | 5346 | 15 | 2019 Dec | 40 | Y | [19] |
| *Alosa sapidissima* | A | 33 | GM | 4354 | 13 | 2019 Dec | 76 | Y | [20] |
| *Alticus arnoldorum* | A | 6 | C | 204 | 17 | 2021 Jan | 8 | N | [21] |
| *Amphiprion bicinctus* | A | 19 | C | 991 | 38 | 2021 Jan | 14 | N | [22] |
| *Amphiprion chrysopterus* | A | 1 | C | 46 | 8 | 2021 Jan | 17 | N | [23] |
| *Amphiprion melanopus* | A | 12 | C | 426 | 22 | 2021 Jan | 12 | N | [24] |
| *Amphiprion sandaracinos* | A | 1 | C | 160 | 8 | 2021 Jan | 14 | N | [23] |
| *Anguilla anguilla* | A | 30 | AC, C, GM | 1336 | 21 | 2019 Dec | 121.5 | Y | [25] |
| *Anguilla rostrata* | A | 40 | AC, C | 2290 | 39 | 2019 Dec | 152 | Y | [25,26] |
| *Atractoscion aequidens* | A | 2 | M | 396 | 8 | 2021 Jan | 130 | Y | [27] |
| *Chaetodon capistratus* | A | 10 | T | 79 | 11 | 2021 Jan | 15 | N | [28] |
| *Chaetodon guttatissimus* | A | 2 | AC, M | 43 | 20 | 2021 Jan | 12 | N | [29] |
| *Chaetodon lunulatus* | A | 7 | AC, M | 263 | 24 | 2021 Jan | 26.7 | N | [30,31] |
| *Chaetodon punctatofasciatus* | A | 2 | AC, M | 25 | 20 | 2021 Jan | 12 | M | [29] |
| *Chaetodon trifascialis* | A | 5 | M | 209 | 11 | 2021 Jan | 18 | N | [30] |
| *Chaetodon trifasciatus* | A | 3 | AC, M | 71 | 12 | 2021 Jan | 15 | M | [31] |
| *Clupea pallasii* | A | 18 | M | 3600 | 5 | 2021 Jan | 46 | Y | [32] |
| *Diplodus vulgaris* | A | 12 | M | 310 | 9 | 2019 Dec | 45 | Y | [33] |
| *Elacatinus lori* | A | 10 | C | 300 | 14 | 2021 Jan | 4.5 | N | [34] |
| *Engraulis encrasicolus* | A | 17 | C, M | 724 | 15 | 2019 Dec | 20 | Y | [35,36] |
| *Fundulus parvipinnis* | A | 23 | C | 182 | 40 | 2021 Jan | 10.8 | N | [37,38] |
| *Gadus morhua* | A | 14 | C | 1330 | 8 | 2019 Dec | 200 | Y | [39] |
| *Gasterosteus aculeatus* | A | 14 | C, AC, M, GM | 545 | 68 | 2019 Dec | 11 | M | [40–44] |
| *Gillichthys mirabilis* | A | 29 | C | 311 | 32 | 2021 Jan | 21 | N | [37,38] |

| | | | | | | | | | |
|---|---|---|---|---|---|---|---|---|---|
| *Gymnosarda unicolor* | A | 6 | M | 73 | 13 | 2021 Jan | 248 | M | 45 |
| *Haemulon flavolineatum* | A | 11 | T | 85 | 11 | 2021 Jan | 30 | Y | 28 |
| *Hypoplectrus nigricans* | A | 9 | T | 66 | 13 | 2021 Jan | 15.2 | N | 28 |
| *Kryptolebias marmoratus* | A | 14 | C | 388 | 65 | 2021 Jan | 7.5 | N | 46,47 |
| *Lethrinus nebulosus* | A | 4 | AC | 350 | 12 | 2021 Jan | 87 | Y | 48 |
| *Limanda limanda* | A | 15 | C | 3006 | 16 | 2019 Dec | 40 | Y | 49 |
| *Lithognathus lithognathus* | A | 3 | C, GM | 50 | 9 | 2021 Jan | 200 | M | 50 |
| *Merluccius capensis* | A | 9 | M, AU | 1477 | 16 | 2021 Jan | 140 | M | 51,52 |
| *Merluccius paradoxus* | A | 9 | M, AU | 1452 | 18 | 2021 Jan | 115 | Y | 51,52 |
| *Naso unicornis* | A | 7 | M | 562 | 8 | 2021 Jan | 70 | Y | 53 |
| *Nerophis lumbriciformis* | A | 2 | C | 155 | 16 | 2019 Dec | 15 | N | 54,55 |
| *Oncorhynchus nerka* | A | 16 | C, GM | 1796 | 36 | 2021 Jan | 84 | Y | 56–58 |
| *Pachymetopon blochii* | A | 2 | C, GM | 50 | 9 | 2021 Jan | 46 | Y | 50 |
| *Paracirrhites arcatus* | A | 7 | C | 264 | 30 | 2021 Jan | 20 | N | 59 |
| *Phycodurus eques* | A | 6 | M | 49 | 7 | 2021 Jan | 35 | N | 60 |
| *Plectropomus leopardus* | A | 8 | C | 1204 | 46 | 2021 Jan | 120 | Y | 61,62 |
| *Plectropomus maculatus* | A | 7 | C | 2477 | 48 | 2021 Jan | 125 | Y | 61,62 |
| *Pomatomus saltatrix* | A | 4 | M | 218 | 15 | 2021 Jan | 130 | Y | 63 |
| *Pomatoschistus flavescens** | A | 3 | C | 83 | 8 | 2019 Dec | 6 | N | 64,65 |
| *Pristipomoides zonatus* | A | 8 | M, GM | 292 | 8 | 2021 Jan | 57.5 | Y | 66 |
| *Quietula y-cauda* | A | 28 | C | 182 | 34 | 2021 Jan | 7 | N | 37,38 |
| *Salmo salar* | A | 281 | C, M, GM | 20038 | 148 | 2019 Dec, 2021 Jan | 150 | Y | 67–77 |
| *Scomberomorus niphonius* | A | 10 | M | 1109 | 5 | 2021 Jan | 113.1 | Y | 78 |
| *Sebastes mentella* | A | 8 | C | 117 | 12 | 2019 Dec | 77.5 | Y | 79 |
| *Serranus cabrilla* | A | 11 | M | 330 | 11 | 2019 Dec | 40 | M | 80 |

| | | | | | | | | | |
|---|---|---|---|---|---|---|---|---|---|
| *Siganus fuscescens* | A | 8 | M | 343 | 12 | 2021 Jan | 40 | Y | 81 |
| *Solea solea* | A | 4 | M | 342 | 11 | 2019 Dec | 70 | Y | 82 |
| *Sparus aurata* | A | 5 | C | 171 | 10 | 2021 Jan | 70 | Y | 83 |
| *Sprattus sprattus* | A | 13 | C, AC | 1285 | 8 | 2019 Dec | 16 | Y | 84 |
| *Stegastes partitus* | A | 15 | T, M | 3471 | 22 | 2021 Jan | 10 | N | 28,85 |
| *Thalassoma bifasciatum* | A | 11 | T | 81 | 11 | 2021 Jan | 25 | N | 28 |
| *Totoaba macdonaldi* | A | 5 | M | 310 | 16 | 2021 Jan | 200 | Y | 86 |

SI Table 2. Model summaries for the effects of human population density and cumulative human impacts on genetic diversity and differentiation across all buffers used (25,50,100 and 200km). Summaries are for models with priors. Mean overall effect sizes for each predictor are given with 95% credible intervals. The widely applicable information criterion (WAIC) is an indicator of model fit. Probability of direction (pd) is a measure of effect existence indicating the certainty for which an effect is positive or negative. $R^2$ values represent coefficient of variation and indicate model fit. Bayesian $R^2$ is the variance of the predicted values divided by the variance of predicted values plus the expected variance of the errors.

| | predictor | mean (95% CI) | pd | Bayesian $R^2$ | WAIC |
|---|---|---|---|---|---|
| gene diversity | human population density (200km) | -0.10 (-0.17 -0.01) | 98.80% | 0.8312344 | 1173.8 |
| | human population density (100km) | -0.08 (-0.16 -0.00) | 97.80% | 0.4079385 | 11675.9 |
| | human population density (50km) | -0.05 (-0.11 0.00) | 96.80% | 0.4161094 | 9223.0 |
| | human population density (25km) | -0.07 (-0.13 -0.00) | 97.79% | NA | NA |
| | cumulative human impacts (200km) | -0.04 (-0.11 0.03) | 88.05% | 0.8065582 | 1374.1 |
| | cumulative human impacts (100km) | -0.03 (-0.09 0.03) | 86.04% | 0.797661 | 1412.1 |
| | cumulative human impacts (50km) | -0.03 (-0.07 0.02) | 87.13% | 0.7949981 | 1419.5 |
| | cumulative human impacts (25km) | -0.03 (-0.08 0.03) | 83.83% | 0.7979978 | 1411.2 |
| allelic richness | human population density (200km) | -0.06 (-0.15 0.02) | 94.23% | 0.7502231 | 1587.1 |
| | human population density (100km) | -0.04 (-0.12 0.05) | 83.70% | 0.3674889 | 7477.5 |
| | human population density (50km) | -0.03 (-0.08 0.03) | 86.31% | 0.3465074 | 6618.1 |
| | human population density (25km) | 0.05 (-0.09 0.19) | 77.22% | NA | NA |
| | cumulative human impacts (200km) | -0.06 (-0.13 0.00) | 97.31% | 0.7464375 | 1661.1 |

| | | | | | |
|---|---|---|---|---|---|
| | cumulative human impacts (100km) | -0.04 (-0.09 0.02) | 91.17% | 0.7368886 | 1690.9 |
| | cumulative human impacts (50km) | -0.00 (-0.05 0.05) | 52.85% | 0.7346569 | 1700.1 |
| | cumulative human impacts (25km) | 0.00 (-0.06 0.05) | 53.99% | 0.7352335 | 1698.1 |
| per population FST | human population density (200km) | 0.04 (-0.12 0.21) | 70.50% | 0.5488618 | 2198.8 |
| | human population density (100km) | -0.01 (-0.19 0.14) | 56.45% | 0.3199093 | 5019.0 |
| | human population density (50km) | -0.02 (-0.13 0.07) | 66.17% | 0.3069676 | 3692.8 |
| | human population density (25km) | -0.02 (-0.13 0.08) | 63.35% | NA | NA |
| | cumulative human impacts (200km) | 0.02 (-0.08 0.12) | 65.75% | 0.488848 | 2408.7 |
| | cumulative human impacts (100km) | -0.01 (-0.08 0.06) | 58.31% | 0.4698496 | 2435.9 |
| | cumulative human impacts (50km) | -0.01 (-0.08 0.05) | 63.19% | 0.4677949 | 2437.4 |
| | cumulative human impacts (25km) | -0.01 (-0.08 0.05) | 61.92% | 0.46809 | 2441.4 |
| effective population size | human population density (200km) | -0.03 (-0.17 0.12) | 67.67% | 0.376307 | 1650.1 |
| | human population density (100km) | -0.09 (-0.23 0.03) | 93.60% | 0.2808449 | 2412.4 |
| | human population density (50km) | 0.00 (-0.08 0.08) | 54.55% | 0.2658967 | 2356.2 |
| | human population density (25km) | -0.03 (-0.13 0.07) | 70.28% | NA | NA |
| | cumulative human impacts (200km) | -0.03 (-0.18 0.12) | 65.35% | 0.3585834 | 1712.7 |
| | cumulative human impacts (100km) | 0.02 (-0.10 0.15) | 63.78% | 0.3391453 | 1726.6 |
| | cumulative human impacts (50km) | 0.02 (-0.09 0.13) | 60.98% | 0.3278958 | 1733.7 |
| | cumulative human impacts (25km) | 0.05 (-0.05 0.17) | 82.38% | 0.3239981 | 1733.7 |

SI Table 3. Model summaries for the effects of human population density and cumulative human impacts on genetic diversity and differentiation across all buffers used (25,50,100 and 200km). Summaries are for models without priors. Mean overall effect sizes for each predictor are given with 95% credible intervals. The widely applicable information criterion (WAIC) is an indicator of model fit. Probability of direction (pd) is a measure of effect existence indicating the certainty for which an effect is positive or negative. $R^2$ values represent coefficient of variation and indicate model fit. Bayesian $R^2$ is the variance of the predicted values divided by the variance of predicted values plus the expected variance of the errors.

| | predictor | mean (95% CI) | pd | Bayesian $R^2$ | WAIC |
|---|---|---|---|---|---|
| gene diversity | human population density (200km) | -0.10 (-0.18 -0.02) | 98.86% | 0.8312535 | 1173.6 |
| | human population density (100km) | -0.08 (-0.16 -0.00) | 97.76% | 0.4080924 | 11708.3 |

| | | | | | |
|---|---|---|---|---|---|
| | human population density (50km) | -0.05 (-0.11 0.00) | 97.15% | 0.4158807 | 9181.0 |
| | human population density (25km) | -0.07 (-0.13 -0.00) | 97.79% | NA | NA |
| | cumulative human impacts (200km) | -0.04 (-0.11 0.03) | 88.19% | 0.8065229 | 1374.3 |
| | cumulative human impacts (100km) | -0.03 (-0.09 0.04) | 85.01% | 0.7975803 | 1412.2 |
| | cumulative human impacts (50km) | -0.03 (-0.07 0.02) | 87.71% | 0.7949353 | 1420.5 |
| | cumulative human impacts (25km) | -0.03 (-0.08 0.03) | 83.67% | 0.7978023 | 1412.8 |
| allelic richness | human population density (200km) | -0.07 (-0.15 0.02) | 94.11% | 0.7503584 | 1585.5 |
| | human population density (100km) | -0.04 (-0.12 0.04) | 85.21% | 0.3677036 | 7469.8 |
| | human population density (50km) | -0.03 (-0.08 0.03) | 85.96% | 0.3464067 | 6625.9 |
| | human population density (25km) | 0.05 (-0.08 0.19) | 78.56% | NA | NA |
| | cumulative human impacts (200km) | -0.06 (-0.12 0.00) | 97.12% | 0.7464818 | 1660.2 |
| | cumulative human impacts (100km) | -0.04 (-0.09 0.02) | 91.66% | 0.7370303 | 1691.0 |
| | cumulative human impacts (50km) | -0.00 (-0.05 0.05) | 53.16% | 0.7346818 | 1698.1 |
| | cumulative human impacts (25km) | 0.00 (-0.05 0.05) | 53.85% | 0.7353002 | 1697.4 |
| per population FST | human population density (200km) | 0.04 (-0.13 0.21) | 68.97% | 0.5486725 | 2199.2 |
| | human population density (100km) | -0.01 (-0.18 0.15) | 56.50% | 0.3193974 | 4999.2 |
| | human population density (50km) | -0.02 (-0.13 0.07) | 66.63% | 0.3075441 | 3695.3 |
| | human population density (25km) | -0.02 (-0.13 0.08) | 62.14% | NA | NA |
| | cumulative human impacts (200km) | 0.02 (-0.08 0.12) | 66.25% | 0.4890242 | 2408.4 |
| | cumulative human impacts (100km) | -0.01 (-0.08 0.06) | 57.48% | 0.4698607 | 2436.7 |
| | cumulative human impacts (50km) | -0.01 (-0.08 0.06) | 62.19% | 0.4677596 | 2436.8 |
| | cumulative human impacts (25km) | -0.01 (-0.08 0.06) | 62.01% | 0.4678947 | 2441.1 |
| effective population size | human population density (200km) | -0.03 (-0.16 0.12) | 68.06% | 0.3773541 | 1649.7 |
| | human population density (100km) | -0.09 (-0.23 0.03) | 93.48% | 0.2813046 | 2419.6 |
| | human population density (50km) | 0.00 (-0.08 0.09) | 53.87% | 0.2659971 | 2344.3 |
| | human population density (25km) | -0.03 (-0.13 0.07) | 71.06% | NA | NA |
| | cumulative human impacts (200km) | -0.03 (-0.17 0.11) | 66.66% | 0.3574458 | 1714.2 |
| | cumulative human impacts (100km) | 0.02 (-0.10 0.15) | 64.04% | 0.3381898 | 1727.6 |
| | cumulative human impacts (50km) | 0.02 (-0.09 0.13) | 61.91% | 0.3268127 | 1734.5 |
| | cumulative human impacts (25km) | 0.05 (-0.05 0.17) | 81.55% | 0.3234502 | 1735.1 |
